# Disrupting SORL1-retromer endosomal recycling in mice drives the age-dependent neurodegenerative signature of Alzheimer’s disease

**DOI:** 10.64898/2026.10.26.761980

**Authors:** Yasir H. Qureshi, Jia Guo, John Rinald, Zongyu Li, Rajesh Kumar Soni, Ananya Govindarajan, Emily Tortora, Genoveva Uzunova, Yanting Yang, Suvarnambiga Kannan, Priyanka Nanayakkara, Carol Troy, Scott A. Small

## Abstract

MRI studies have mapped the atrophic signature that distinguishes Alzheimer’s disease (AD), isolating regions differentially vulnerable to its age-dependent neurodegenerative process. We test the hypothesis that an endosomal recycling pathway regulated by the interaction of SORL1 with retromer, which is pathogenically disrupted in AD, might drive the neurodegenerative process via multiple yearlong studies in over 140 wildtype and SORL1 deficient mice, modeling AD’s causative mutations. MRI studies revealed that SORL1 deficiency induced an age-dependent AD-pattern of atrophy, centered on the trans-entorhinal cortex, and that atrophy was prevented by SORL1 restoration, secondary to overexpressing the retromer protein VPS26b. Postmortem studies validated that the atrophy reflects neuronal loss. Proteomic profiling revealed that the atrophy is associated with reductions in cell death suppressors, one of which was then found deficient in the trans-entorhinal cortex of AD patients. Besides supporting the hypothesis, taken together with other findings, the results suggest that SORL1-retromer disruptions represent a therapeutically-amenable common ‘neurodegeneration pathway’ in AD.

## INTRODUCTION

Neurodegenerative disorders manifest with unique neuroanatomical signatures of MRI-detected atrophy^1^, reflecting regional sensitivities to each disorder’s distinct neurodegenerative process. MRI studies have established that the atrophic signature that distinguishes Alzheimer’s disease (AD) is predominately centered on the trans-entorhinal cortex^2–5^, supported by evidence of hypometabolism^6^ and neuronal loss^7,8^. As indicated by mapping atrophy over time^9^, other cortical regions are secondarily sensitive to the neurodegenerative process, but the trans-entorhinal cortex is found reliably atrophic in AD’s preclinical stage^3^ and even during later dementia stages the atrophy there is most pronounced^5^. Collectively, these and related studies justify why this region has been called AD’s “ground zero”^10^.

One biological pathway hypothesized to trigger AD’s neurodegenerative process is an endosomal recycling pathway that is regulated by the interaction of the sortilin receptor SORL1 with the retromer trafficking complex, a heterotrimer comprised of VPS26, VPS35, and VPS29^11^. Based on an integration of genetics with studies in human neurons, animal models, and patient brains, pathway disruptions are implicated as common and causally pathogenic in AD^12^. The pathway is normally dedicated to synaptic plasticity by orchestrating many cellular factors, including the recycling of receptors and the sorting of proteins that control lysosome, autophagosome, and mitochondrial function^13–27^. This can explain why neurons are enriched in SORL1^28–32^ and differentially express the paralogue VPS26b, which is dedicated to neuronal endosomal recycling^5^. It is to VPS26b that SORL1 binds at endosomal membranes, establishing a neuronal SORL1-retromer functional unit^12^.

Many of these factors can be considered neuroprotective^20^ supporting the hypothesis that, in conditions that pathologically disrupt the pathway, it can switch from one that normally supports synaptic plasticity to one that can lead to neuronal demise. Empirical evidence is starting to support the hypothesis, but informatively, even with complete retromer depletion, studies in animal models suggest that neuronal loss occurs in a slowly progressive, “neurodegenerative”, manner^33,34^. This slowness likely reflects the cellular dynamics of how even arresting the pathway is not equivalent to an immediate and widespread depletion of the pathway’s cargo, and there are potentially compensatory alternative recycling pathways^35^. Nevertheless, showing that pathway disruptions induce an age-dependent AD-like pattern of atrophy centered on the trans-entorhinal cortex, and that pathway corrections prevent this atrophy, will mechanistically validate the hypothesis.

SORL1 deficient mouse models are best suited to mechanistically test the hypothesis, since SORL1 haploinsufficiency caused by truncation mutations are, based on genetic epidemiology, considered as causally pathogenic as mutations in the amyloid precursor protein (APP) and the presenilins^36^. Accordingly, we investigated a previously characterized SORL1 deficient mouse model that is genetically engineered to express a targeted deletion of the 5’ region of SORL1 exon4^37^. While these mice express low levels of exon4-deleted SORL1 protein^38^, and it is difficult to discriminate the exon4-deleted SORL1 from wildtype full-length SORL1 protein, an examination of the brains of both SORL1^Exon4+/-^ and SORL1^Exon^^4-/-^ mice showed that the deletion results in a gene-dose dependent reduction in SORL1 levels and function^38^.

As established in patients by imaging the entire brain *in vivo* and over time, MRI-detected regional atrophy has emerged as the most reliable readout to precisely map the atrophic signatures of different neurodegenerative processes^1^. To test whether SORL1 deficiency might induce AD’s atrophic pattern, we therefore rely on a dedicated small-bore 9.4 Telsa (T) scanner equipped with a cryogenically cooled mouse brain RF coil that increases signal-to-noise, important for imaging the small mouse brain, and utilized acquisition and post-acquisition protocols optimized to map mouse brain atrophy^39^. Beginning with MRI and extending to a series of *ex vivo* studies, we have completed multiple yearlong studies in over 140 SORL1 deficient and wildtype mice. In so doing, we were able to show that SORL1 deficiency induces the AD pattern of cortical atrophy, that VPS26b overexpression and SORL1 restoration prevent the atrophy, and that MRI-detected atrophy reflects bona fide neurodegeneration. Moreover, we identify potential neurodegenerative mechanisms via proteomics, which we then find occurs in the atrophic trans-entorhinal cortex of patients with sporadic AD.

Besides supporting the hypothesis, we discuss the details of our findings in the context of previous observations that offer mechanistic insight into why the neurodegenerative process is age-dependent and chronically progressive, a central unresolved question in the field, one that can generalize to the common form of AD. Additionally, we discuss how the results justify developing interventions designed to enhance SORL1-retromer, directly targeting the neurodegenerative process in AD.

## RESULTS

### Mapping the spatiotemporal atrophic pattern in SORL1 deficient mice

In anticipation of a large-scale longitudinal MRI study, we first completed an exploratory study imaging six 12-month-old SORL1^Exon4+/-^ and six SORL1^Exon4+/+^ wildtype mice to determine whether there is any evidence of volume loss and to inform as to the estimated number of mice needed. A voxel-based analysis was performed across the cortex, comparing SORL1^Exon4+/-^ to SORL1^Exon4+/+^ mice, revealing volume loss in the posterior brain, and after correcting for multiple comparisons, the most significant volume reduction is centered in the trans-entorhinal cortex (TEC) (Fig. 1), which is a border zone region between the lateral entorhinal cortex and the perirhinal cortex^5^.

**Figure 1.**
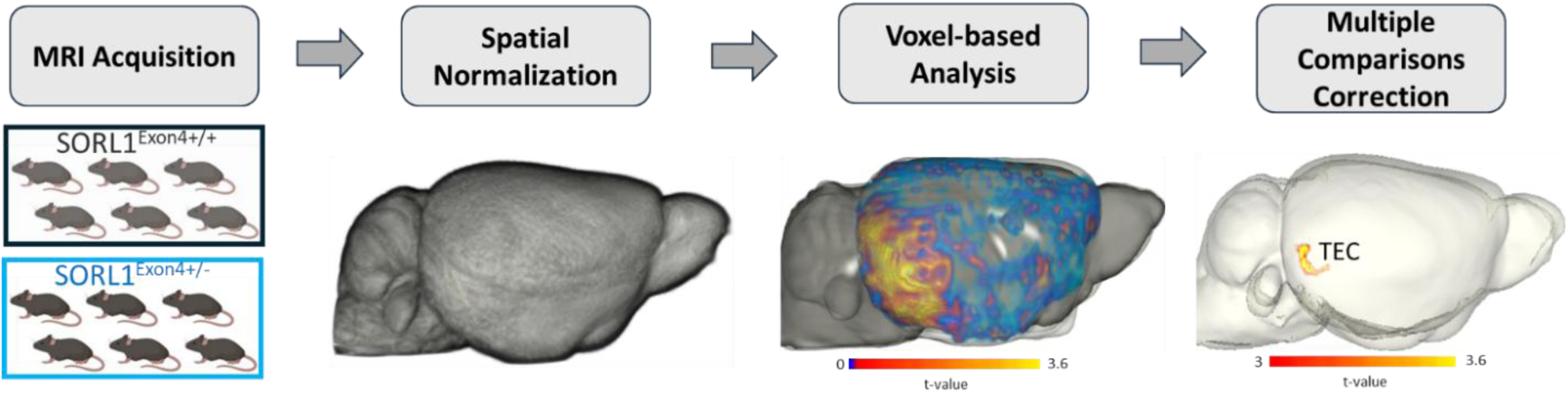
Exploratory MRI study and post-acquisition processing stream. At 12 months, SORL1^Exon4+/-^ mice (n = 6) and age-matched wildtype controls (n = 6; non-littermates) underwent structural MRI with a 9.4 T Bruker BioSpec scanner equipped with a CryoProbe. Images were spatially normalized for tensor-based morphometry (TBM) volumetric analysis. Voxel-wise comparison of the groups identified cortical regions with reduced volume. When corrected for multiple comparisons, the most reliable volume reduction localized to the trans-entorhinal cortex (TEC). The corrected map used a voxel-wise cluster-forming threshold of *P* < 0.005 and cluster-wise corrected *P* < 0.05.

Informed by this exploratory study, we imaged fourteen SORL1^Exon4+/-^ mice, nine SORL1^Exon^^4-/-^ mice, and ten SORL1^Exon4+/+^ littermates at 3 months of age, all with an equal mix of females and males, and reimaged them at 12 months of age. We also imaged a small subset of these longitudinally-followed mice at 6 months of age, and we supplemented this age with additional mice, for a total of nine SORL1^Exon4+/-^ mice, eight SORL1^Exon^^4-/-^ mice, and eight SORL1^Exon4+/+^ wildtype littermates.

A voxel-based linear regression model was applied to all 12 month old mice across genotypes showing volume reductions in posterior and frontal brain regions (Fig 2B). As in the exploratory study, the multiple comparisons corrected map shows that the posterior volume reductions centered in the TEC (Fig. 2B). In this larger study using both SORL1 deficient models, the frontal cortex reductions were centered in the anterior prefrontal cortex (Fig. 2B). By repeating the analysis including sex as a covariate, no significant sex differences were detected in the vulnerability maps.

**Figure 2.**
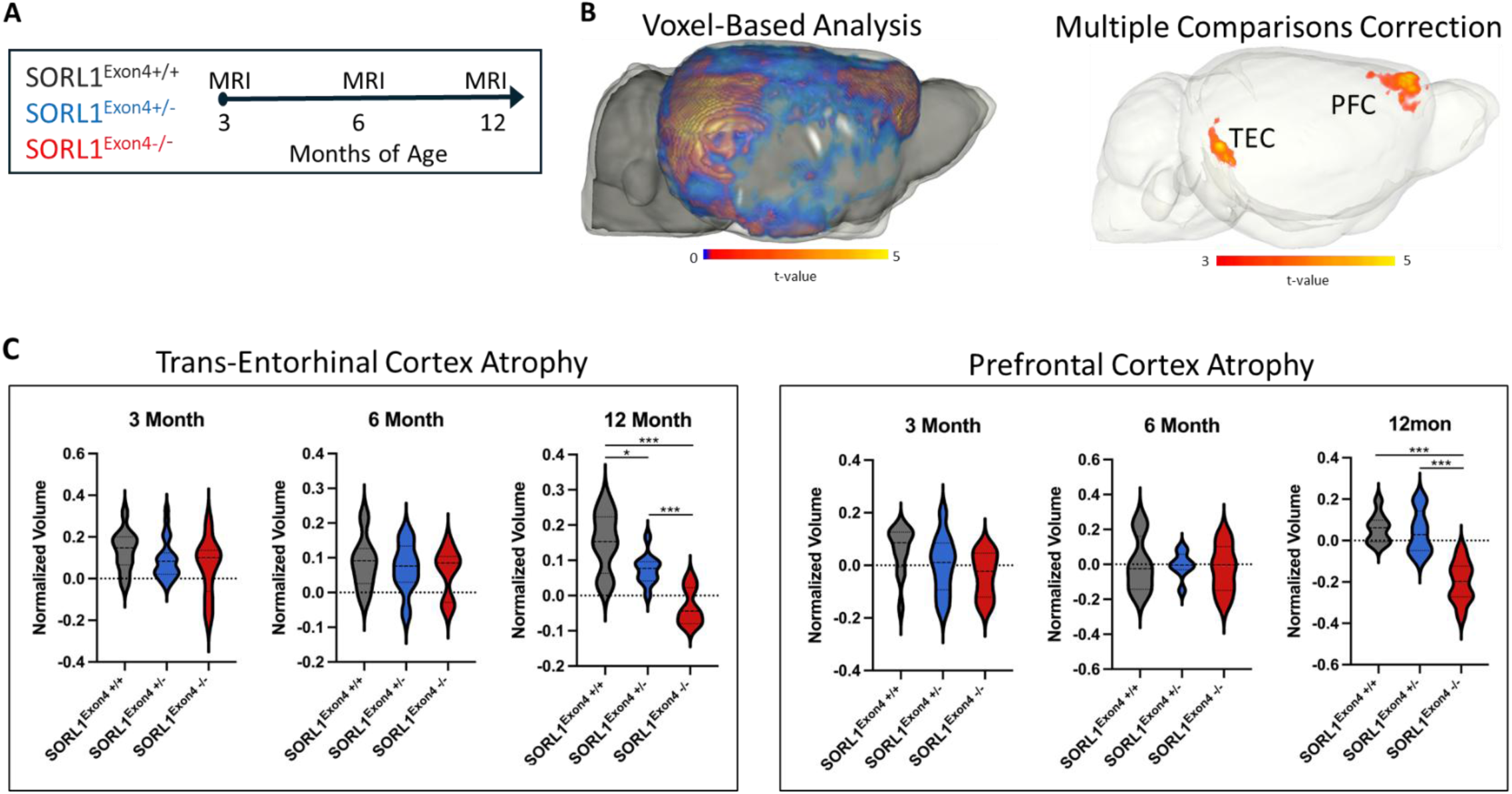
SORL1 deficiency phenocopies the cortical atrophic signature of late-onset Alzheimer’s disease. (A) Genotype cohorts were studied at 3 and 12 months, including animals rescanned longitudinally when available. The intermediate, approximately 6-month population included both repeat scans and additional age-matched animals. MRI scan counts by genotype and sex (M, male; F, female) were: 3-month SORL1^Exon4+/-^ mice (n = 14; M = 7, F = 7), SORL1^Exon^^4-/-^ mice (n = 9; M = 4, F = 5), and SORL1^Exon4+/+^ wildtype littermates (n = 10; M = 4, F = 6); 6-month SORL1^Exon4+/-^ mice (n = 9; M = 6, F = 3), SORL1^Exon^^4-/-^ mice (n = 8; M = 2, F = 6), and SORL1^Exon4+/+^ wildtype mice (n = 8; M = 4, F = 4); and 12-month SORL1^Exon4+/-^ mice (n = 14; M = 7, F = 7), SORL1^Exon^^4-/-^ mice (n = 9; M = 4, F = 5), and SORL1^Exon4+/+^ wildtype littermates (n = 10; M = 4, F = 6). (B) To identify regions-of-interest, a linear regression was applied to all 3 genotypes at 12 months of age, and the left image shows the resultant voxel-based analysis mapping a gene-dose dependent effect. When corrected for multiple comparisons, as shown in the right image, 2 regions showed the most reliable volume reductions, the trans-entorhinal cortex and prefrontal cortex. The corrected 12-month log-JD analysis used a voxel-wise cluster-forming threshold of *P* < 0.005 and cluster-wise *P* < 0.05; the resulting TEC and PFC clusters defined the ROIs used for the regional volume measurements in panel C. (C) A TEC region-of-interest was applied to the whole data set and, as shown in the left graph, gene-dose dependent atrophy emerged at 12 months of age. A PFC region-of-interest was applied to the whole data set and, as shown in the right graph, atrophy also emerged at 12 months of age, but only in the SORL1^Exon^^4-/-^ mice. Regional volume was calculated for each scan by summing JD values within the corresponding ROI; the plots show normalized regional volumes for the age-specific MRI populations described in panel A. *\*P < 0.05, **P < 0.01, ***P < 0.001*.

We accordingly derived a TEC and a prefrontal cortex region-of-interest and applied them to the full longitudinal study. As shown in Figure 2B, TEC atrophy was found to emerge only at 12 months with a gene-dose dependent worsening of atrophy. Prefrontal cortex atrophy also emerged at 12 months, but only in the SORL1^Exon^^4-/-^ mice, not in the SORL1^Exon4+/-^ mice.

Collectively, the atrophic patterns show that the TEC is most sensitive, and that higher order cortical regions are secondarily sensitive, to SORL1 deficiency, generally overlapping with the differential atrophic sensitivities observed in AD^5^. Since the atrophic pattern emerges in midlife, SORL1 deficiency appears to phenocopy AD’s TEC-centered spatiotemporal pattern of MRI-detected cortical atrophy.

### SORL1 deficiency worsens with age

To investigate the relationship between SORL1 levels and atrophy, we microscopically dissected the TEC in cohorts of all mouse genotypes at 3 and 12 months of age and dissected the frontal cortex (FC) in the 12 month cohort, and samples were immunoblotted for SORL1.

SORL1 levels observed across all genotypes at 3 months of age showed a gene-dose dependent pattern and degree of SORL1 deficiency (Fig. 3A) that is nearly identical to that previously described at an equivalent age^38^. Interestingly, at 12 months of age, we find the same gene-dose dependent pattern, but now the SORL1 levels show a greater reduction (Fig. 3A). An examination of the patterns suggested that atrophy is associated with >50% SORL1 deficiency. We further confirmed this interpretation by showing a >50% SORL1 deficiency in the FC of 12-month-old SORL1^Exon^^4-/-^ mice (Fig. 3B).

**Figure 3.**
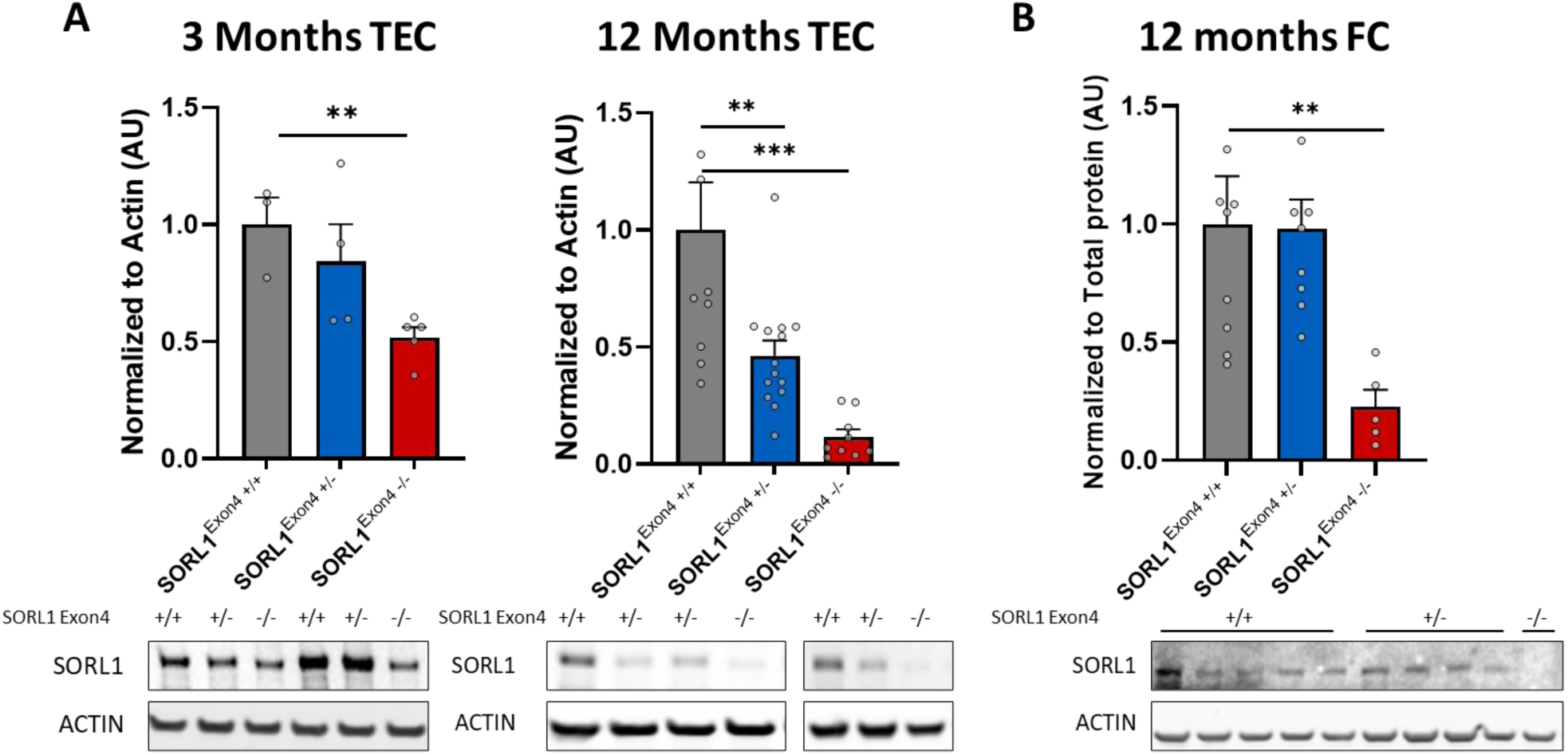
SORL1 levels are associated with regional atrophy. (A) The trans-entorhinal cortex shows a gene-dose dependent pattern of SORL1 deficiency that worsened with age, suggesting that a reduction in SORL1 levels greater than 50% is associated with atrophy (n = 3, 4, & 5 at 3 months and 8, 14, & 9 at 12 months). (B) SORL1 deficiency greater than 50% is found in the frontal cortex at 12 months, but only in SORL1^Exon^^4-/-^ mice (n = 8, 8, & 5). Data are mean ± SEM and genotypes are compared by two-tailed unpaired Student’s t-test. *\*P < 0.05, **P < 0.01, ***P < 0.001*.

Together, the results suggest that the atrophic process occurs in association with a critical threshold of reduced SORL1 levels that emerges in an age-dependent manner, at a level that is similar to the SORL1 reductions observed in the atrophic TEC of sporadic AD patients^5^.

### VPS26b overexpression restores SORL1 levels and prevents atrophy

VPS26b is the retromer protein that directly binds SORL1, establishing a neuronal SORL1-retromer functional unit^12^. To mechanistically confirm that the atrophy observed in SORL1 deficient mice is driven by impairments in SORL1-retromer endosomal recycling we tested the effect of overexpression of VPS26b, using an AAV9-VPS26b protocol whose route of delivery results in reliable overexpression in the entorhinal cortex and surrounding medial temporal lobe regions^5^. Besides the large size of SORL1, one of the largest in the genome which prohibits its packaging into viral vectors, the reliance on the neuronal selective VPS26b will ensure that any effects are localized primarily to neurons and linked to SORL1-retromer endosomal recycling.

Accordingly, a third cohort of SORL1^Exon4+/-^ and SORL1^Exon4+/+^ mice were generated, and AAV9-VPS26b was delivered to ten 3-month-old SORL1^Exon4+/-^ mice and an AAV9-empty vector control was delivered to eight 3-month-old SORL1^Exon4+/-^ mice. At 12 months of age, MRI was performed in AAV9-VPS26b-treated SORL1^Exon4+/-^ mice, AAV9-empty-vector-treated SORL1^Exon4+/-^ mice, and SORL1^Exon4+/+^ wildtype mice. As shown in Figure 4B, compared to SORL1^Exon4+/+^ mice, treatment of SORL1^Exon4+/-^ mice with AAV9-VPS26b prevented the onset of atrophy. By comparing 12-month-old SORL1^Exon4+/-^ mice who received the empty vector control to SORL1^Exon4+/+^ wildtype mice, we replicated for a third time that SORL1 deficiency reliably results in TEC atrophy.

**Figure 4.**
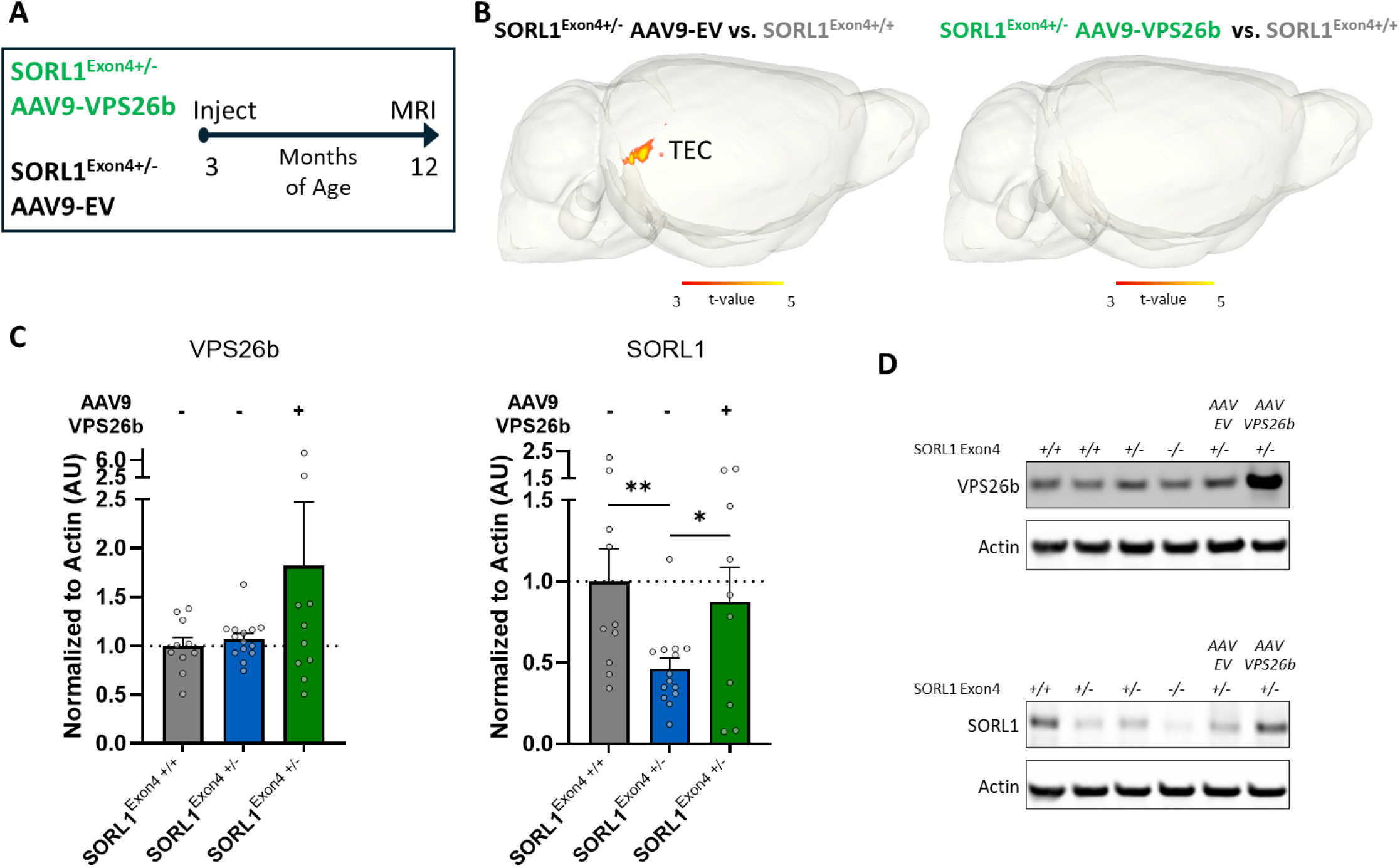
VPS26b overexpression prevents atrophy and restored SORL1 deficiency. (A) At 3 months, SORL1^Exon4+/-^ mice received AAV9-VPS26b (n = 10 injected) or AAV9-empty vector (EV) (n = 8 injected). At 12 months, the MRI comparison included available VPS26b-treated SORL1^Exon4+/-^ mice (n = 7; M = 4, F = 3), empty-vector SORL1^Exon4+/-^ mice (n = 8; M = 4, F = 4), and SORL1^Exon4+/+^ wildtype mice (n = 9; M = 5, F = 4). (B) Using the 12-month MRI groups specified in panel A, as shown in the left image, trans-entorhinal cortex atrophy was the most reliable finding when comparing 12-month-old SORL1^Exon4+/-^ mice who received the empty vector control to SORL1^Exon4+/+^ wildtype mice. As shown in the right image, compared to SORL1^Exon4+/+^ wildtype mice, treatment of SORL1^Exon4+/-^ mice with AAV9-VPS26b prevented the onset of atrophy. The voxel-based log-JD comparisons used a voxel-wise cluster-forming threshold of *P* < 0.005 and cluster-wise *P* < 0.05. (C & D) 12-month-old trans-entorhinal cortex tissue from the treated mice confirmed that the treatment results in increased VPS26b expression. VPS26b overexpression was associated with increased SORL1 levels, effectively restoring SORL1 deficiency. Data are mean ± SEM and genotypes are compared by two-tailed unpaired Student’s t-test. (n = 10, 14, & 10) *\*P < 0.05, **P < 0.01*.

Immunoblotting microdissected 12-month-old TEC tissue from the treated mice showed a numerical increase in SORL1 by AAV9-EV (Supplementary Fig. 1). We therefore compared the AAV9-VPS26B treated mice with the untreated SORL1^Exon4+/-^ mice, and as expected identified an increase in VPS26b expression. More importantly, VPS26b overexpression led to increased SORL1 levels, effectively restoring SORL1 deficiency (Figure 4C&D).

Collectively, the remarkable rescue of atrophy associated with VPS26b-dependent SORL1 restoration mechanistically confirms that the atrophy is driven by disruptions to neuronal SORL1-retromer endosomal recycling.

### Trans-entorhinal cortex atrophy reflects synaptic and neuronal loss

To confirm that the MRI-detected atrophy reflects the neurodegenerative process, at the end of the MRI study we tested for evidence of synaptic loss. We immunoblotted microdissected TEC tissue for synaptophysin, in mice that were approximately 15 months of age and observed a gene-dose dependent loss of synaptophysin (Fig. 5B&C).

**Figure 5.**
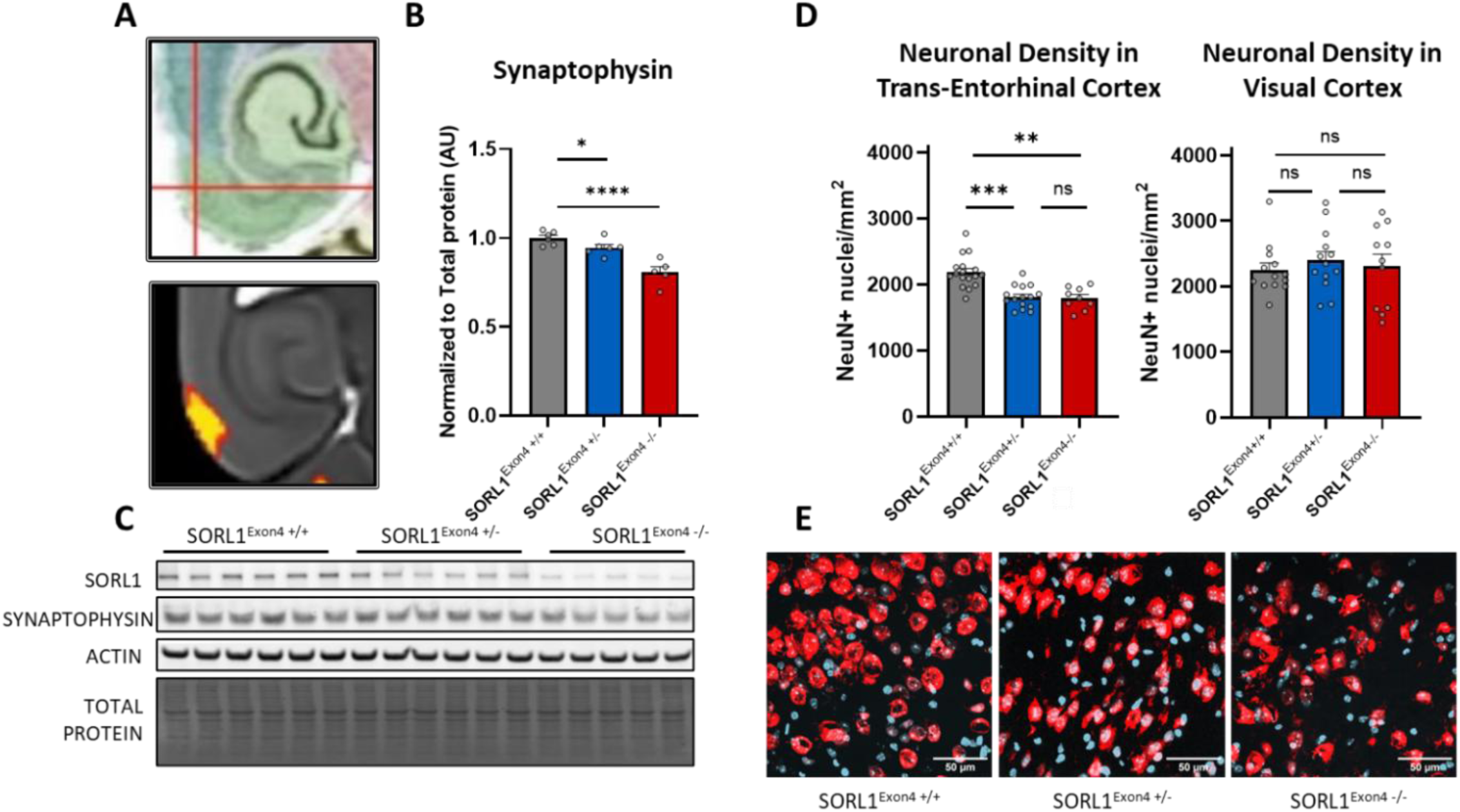
Atrophy is associated with postmortem markers of neurodegeneration. (A) The coordinates for the region with significant volume reduction by MRI. This area was used in the analysis shown in B-E. (B & C) There is gene-dose dependent loss of synaptophysin protein in trans-entorhinal cortex tissue at approximately 15 months of age measured by western blotting. Data are mean ± SEM and genotypes are compared by two-tailed unpaired Student’s t-test. (n = 6, 6, & 5) *\*P < 0.05, ****P < 0.0001.* (D) There is a gene-dose dependent reduction in trans-entorhinal cortex NeuN staining but not in the visual cortex, measured by immunohistochemistry and confocal imaging. The bar graph shows the quantification of NeuN+ nuclei per mm^2^ of cortex. Data are mean ± SEM and genotypes are compared by Kruskal-Wallis test. *\*P < 0.05, **P < 0.01; ***P < 0.001, ns = not significant.* (E) Representative immunofluorescent images of NeuN (red) and DAPI (blue) stained in the trans-entorhinal cortex of aged SORL1 deficient mice (n=3-4 mice per genotype). Scale bar, 50µm.

Bona fide neurodegeneration is characterized by neuronal loss. We therefore generated and aged a fourth cohort of SORL1 deficient mice to approximately 15 months of age and used confocal microscopy to map the neuronal marker NeuN in brain slices that contained the TEC and neighboring regions and observed reduction in TEC NeuN staining (Fig. 5D&E). In addition, we found no changes in NeuN in the visual cortex (Fig. 5D&E). We then aged a fifth cohort, this time testing for neuronal loss at 9 months of age, showing that at this age group no neuronal loss was detected (Supplementary Fig. 2)

Together, the results confirm that the MRI-detected TEC atrophy is reflective of bona fide neurodegeneration, including age-dependent neuronal loss to a degree that phenocopies the level of neuronal loss observed in the entorhinal cortex of early-stage disease^7,8^.

### Trans-entorhinal cortex proteomics suggests neurodegenerative mechanisms

To generate mechanistic clues that might explain why the TEC shows neurodegeneration in the older mice, we generated and aged a sixth cohort of SORL1 deficient mice and via Liquid Chromatography-Tandem Mass Spectrometry (LC-MS/MS) completed a proteomic screen of the TEC microdissected from seven 15-month SORL1^Exon4+/-^ mice and from eight SORL1^Exon4+/+^ littermates. All analyses were performed on FDR corrected t-tests comparing individual protein abundance of SORL1 deficient to wildtype mice. Quality control assessment was performed by PCA and one sample from each group were identified as technical outliers and were excluded prior to differential abundance testing.

First, gene ontology analysis of the whole FDR corrected proteomic dataset identified four enriched ‘biological pathways’ (Fig. 6A): ‘Positive regulation of protein localization to endosome’(GO:1905668); ‘positive regulation of protein localization to early endosomes’ (GO:1902966); ‘protein retention in Golgi apparatus’ (GO:0045053); ‘regulation of adipose tissue development’ (GO:1904177); ‘negative regulation of triglyceride metabolic process’ (GO: 0090209). The first three validate that SORL1 deficiency results in an endosomal phenotype, and the last two reveal a link between SORL1 and lipid metabolism, which has previously been identified in transcriptomic profiling of SORL1 deficient human IPSC-derived neurons^18^.

**Figure 6.**
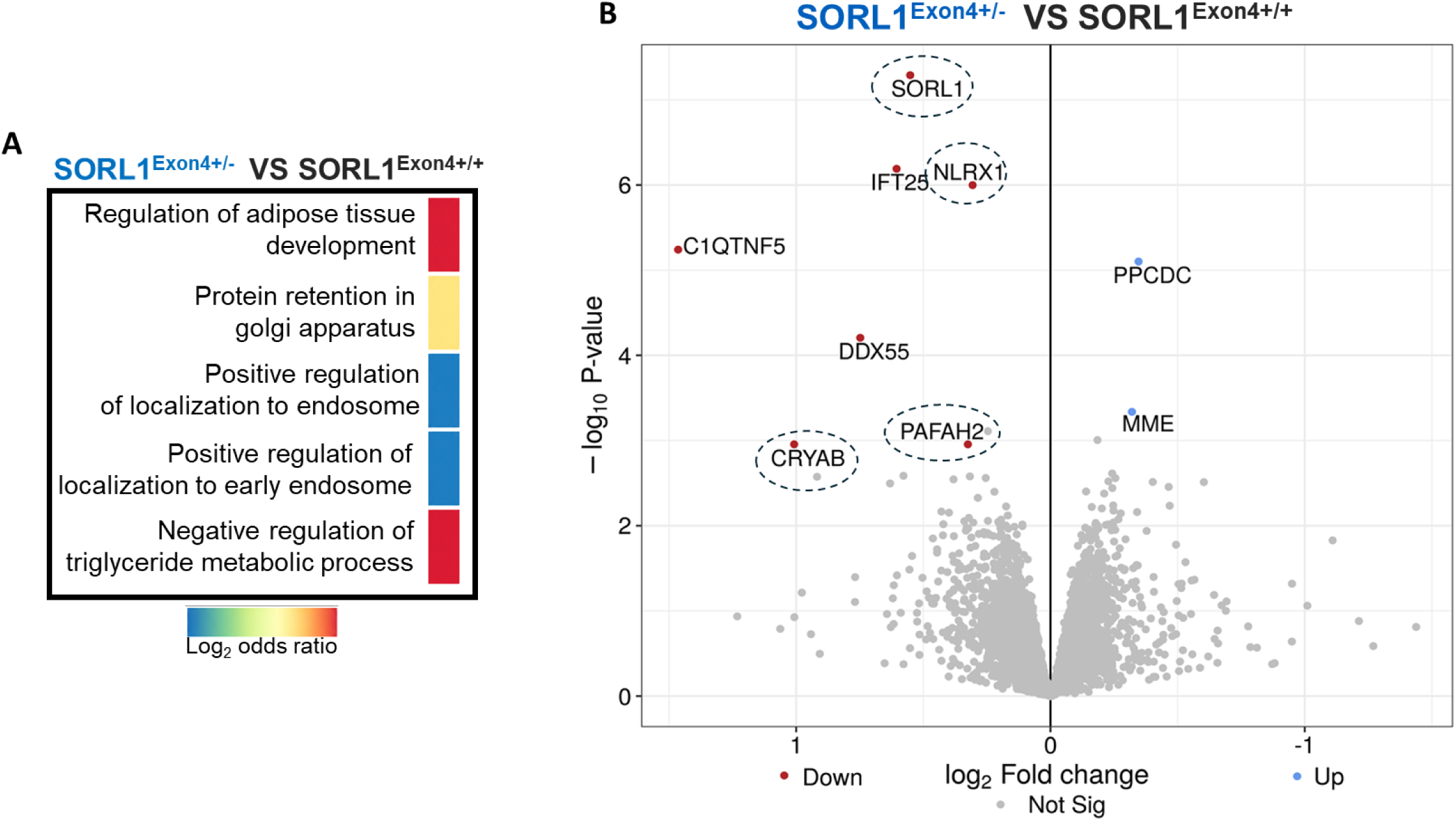
Proteomic analysis in 15 month old SORL1 deficient mice. (A) Pathway Enrichment was performed in DIA-Analyst on all significantly regulated proteins comparing SORL1^Exon4+/-^ vs SORL1^Exon4+/+^ mice. (B) Volcano plot of bulk label-free quantitative proteomics (in-solution digestion) of TEC brain tissue from 15-month-old SORL1^Exon4+/−^ mice and wild-type SORL1^Exon4+/+^ littermates (n = 6-7 per genotype). Log_2_-transformed, normalized protein intensities were analyzed in DIA-Analyst using a moderated t-test, and P values were corrected for multiple testing using t-statistic–based false discovery rate (FDR) correction. The x axis shows log_2_ fold change (SORL1^Exon4+/−^ vs. SORL1^Exon4+/+^), and the y axis shows the unadjusted −log_10_ P value. Proteins with an FDR-adjusted P < 0.05 and |log_2_ fold change| > 0.25 are labeled and colored red (decreased) or blue (increased); all other quantified proteins are shown in gray.

Next, a differential abundance analysis was performed using an FDR-corrected t-test to compare SORL1 deficient mice to wild-type littermates. Significantly altered proteins are displayed in a volcano plot (Fig. 6B), plotting log_2_ fold change against log_2_ adjusted p-value for each protein. SORL1 deficiency at a level that agrees with immunoblotting was observed as a significantly altered protein, acting as a positive control and validating the experimental paradigm. Among the handful of reductions that emerged as significantly altered, we note that NLRX1 (NLR family member X1, also known as NOD5 or NOD9) is a mitochondrial NOD-like receptor (Nucleotide-binding oligomerization domain-like receptor) that can suppress cell death by enhancing mitophagy and attenuating mitochondrial ROS accumulation^40,41^; CRYAB (αB-crystallin) is a suppressor of apoptosis by inhibiting Bax (BCL2 Associated X, Apoptosis Regulator) translocation to mitochondria and blocking downstream caspase-3 activation^42,43^; PAFAH2 (platelet-activating factor acetylhydrolase 2) is a suppressor of ferroptosis through its phospholipase activity that detoxifies phospholipids^44,45^.

To confirm and potentially extend these results we aged a seventh cohort of SORL1 deficiency mice and performed a second proteomic study but this time profiling the TEC of four SORL1^Exon4+/-^ mice and five SORL1^Exon4+/+^ littermates at 9 months, an age that antedates at beginning of the neurodegenerative process (Supplementary Fig. 3). When completing the same analysis as for the 15-month cohort, either because a younger or a smaller dataset, no proteins met the statistical threshold as significantly altered. Nevertheless, SORL1, NLRX1, and CRYAB were found among the differentially reduced proteins in SORL1 deficient mice (Fig. 7A). To investigate how these and PAFAH2 changed from 9 to 15 months of age, we used a multifactorial ANOVA, where age (9 vs.15 months) and genotype (SORL1^Exon4+/+^ vs. SORL1^Exon4+/-^) were included as the fixed factor, and the abundance of each protein was included as the dependent variables. As illustrated in Fig. 7A, the results revealed that, compared to SORL1^Exon4+/+^ littermates, protein reductions for SORL1, CRYAB, and NLRX1 were similar at both age-groups in the SORL1 deficient mice, acting to confirm the reliability of these findings. For PAFAH1, the reduction was not found at 9 months, and for which a significant age X genotype interaction was observed.

**Figure 7.**
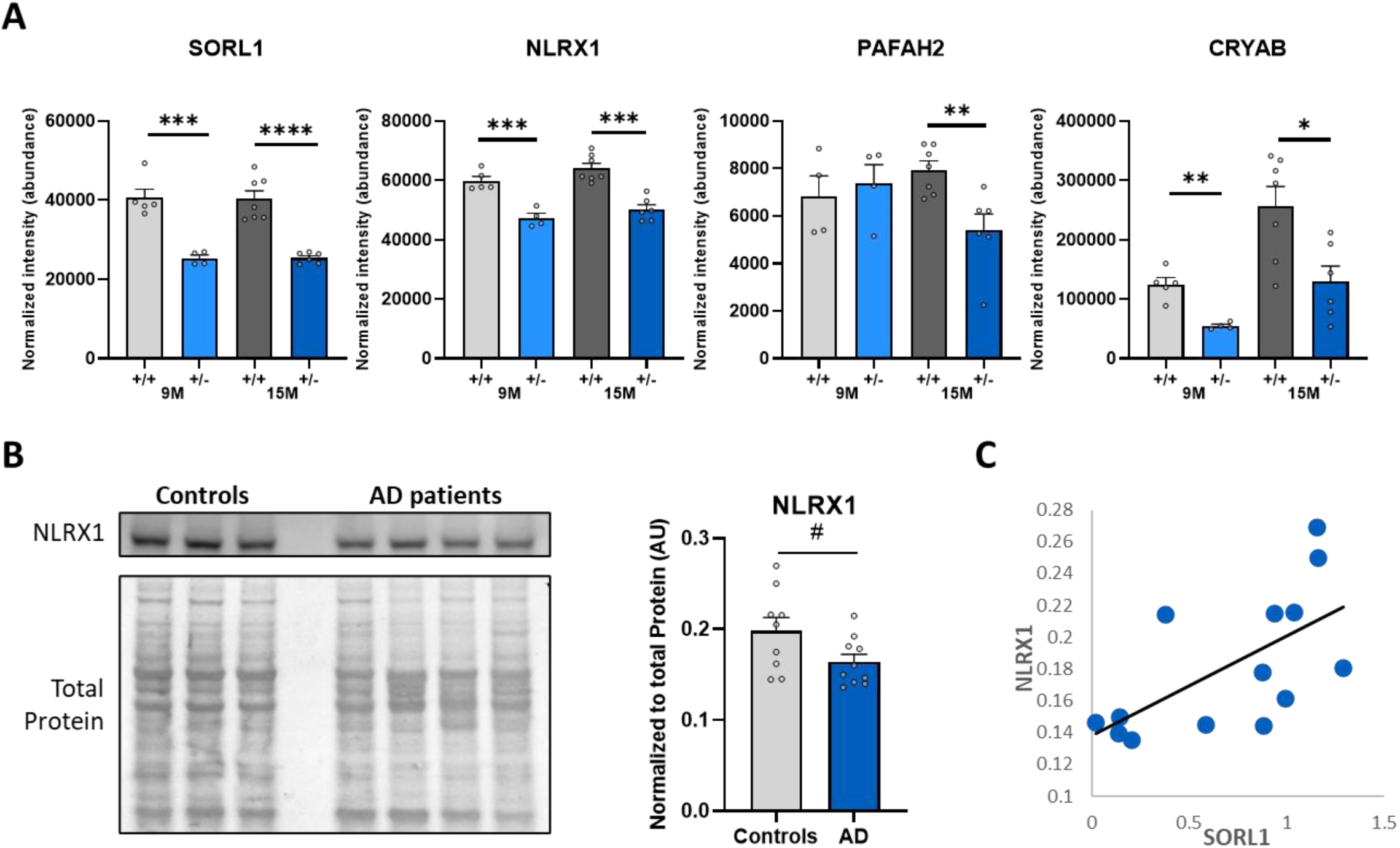
Protein analysis in SORL1 deficient mice and in Alzheimer’s brains. **(A)** Normalized protein intensities of SORL1, NLRX1, PAFAH2 and CRYAB in SORL1^Exon4+/+^ and SORL1^Exon4+/−^ mice at 9 months (9M; n = 4-5 per genotype) and 15 months (15M; 6-7 per genotype) of age. Each dot represents one mouse. Genotypes were compared within each age by two-tailed unpaired Student’s t-test; unannotated comparisons were not significant. **(B)** Representative immunoblots of NLRX1 in postmortem trans-entorhinal cortex (TEC) lysates from healthy controls and patients with Alzheimer’s disease (AD). Total protein staining served as the loading control. Quantification of NLRX1 normalized to total protein (arbitrary units, AU) in healthy controls (n = 9) and patients with AD (n = 10). Each dot represents one individual. P = 0.056, two-tailed unpaired Student’s t-test. **(C)** Correlation between SORL1 and NLRX1 protein levels in the TEC of individuals from “B” for whom SORL1 measurements were available (n = 14; 7 healthy controls and 7 AD). SORL1 levels were determined by immunoblotting and previously reported (Simoes et al., 2021); NLRX1 values are those shown in “B”. Pearson’s r = 0.63 (R^2^ = 0.40), two-tailed P = 0.016; Spearman’s ρ = 0.71, two-tailed P = 0.0045. Bar graphs show mean ± SEM. *\*P < 0.05, **P < 0.01, ***P < 0.001, ****P < 0.0001*.

Finally, we explore whether these findings might be observed in the atrophic TEC of sporadic AD brains. Among the hits, NLRX1 was statistically most reliable, and so we probed samples from the atrophic TEC of sporadic patients and age-matched controls, in which we previously identified SORL1 and VPS26b deficiency^5^. Remarkably, we find an approximately 20% NRLX1 reduction, matching the reduction observed in the TEC of SORL1 deficient mice (Fig. 7B) and that SORL1 levels correlated with NRLX1 (Fig. 7C).

The most informative findings that relate to neurodegeneration is that the neuronal loss observed in SORL1 deficient mice is associated with loss of neuroprotection and cell death suppression, a mechanism that we were able to find in the atrophic TEC of sporadic AD patients.

### Markers of endosomal recycling in trans-entorhinal cortex and other brain regions

Trans-entorhinal cortex neurons are functionally distinguished by receiving a persistent convergence of synaptic stimulation from across the cortex^46^, a network property that can explain why this neuronal population is found to differentially engage in, and to be differentially reliant on, SORL1-retromer endosomal recycling^5,47^. From 6-month-old C57BL/6J wildtype mice, we therefore microdissected the TEC and frontal cortex, as well as other regions of the medial temporal lobe -- the entorhinal cortex, the perirhinal cortex, and the hippocampal CA1 region. We immunoblotted for specific markers of the endosomal recycling pathway, SNX27 (sorting nexin 27) and syntaxin13.

As shown in Figure 8, compared to other regions, the most reliable differential TEC enrichment was found for the retromer recycling-associated Syntaxin13. SNX27 also showed the highest expression in the TEC but was only reliably different from the frontal cortex.

**Figure 8.**
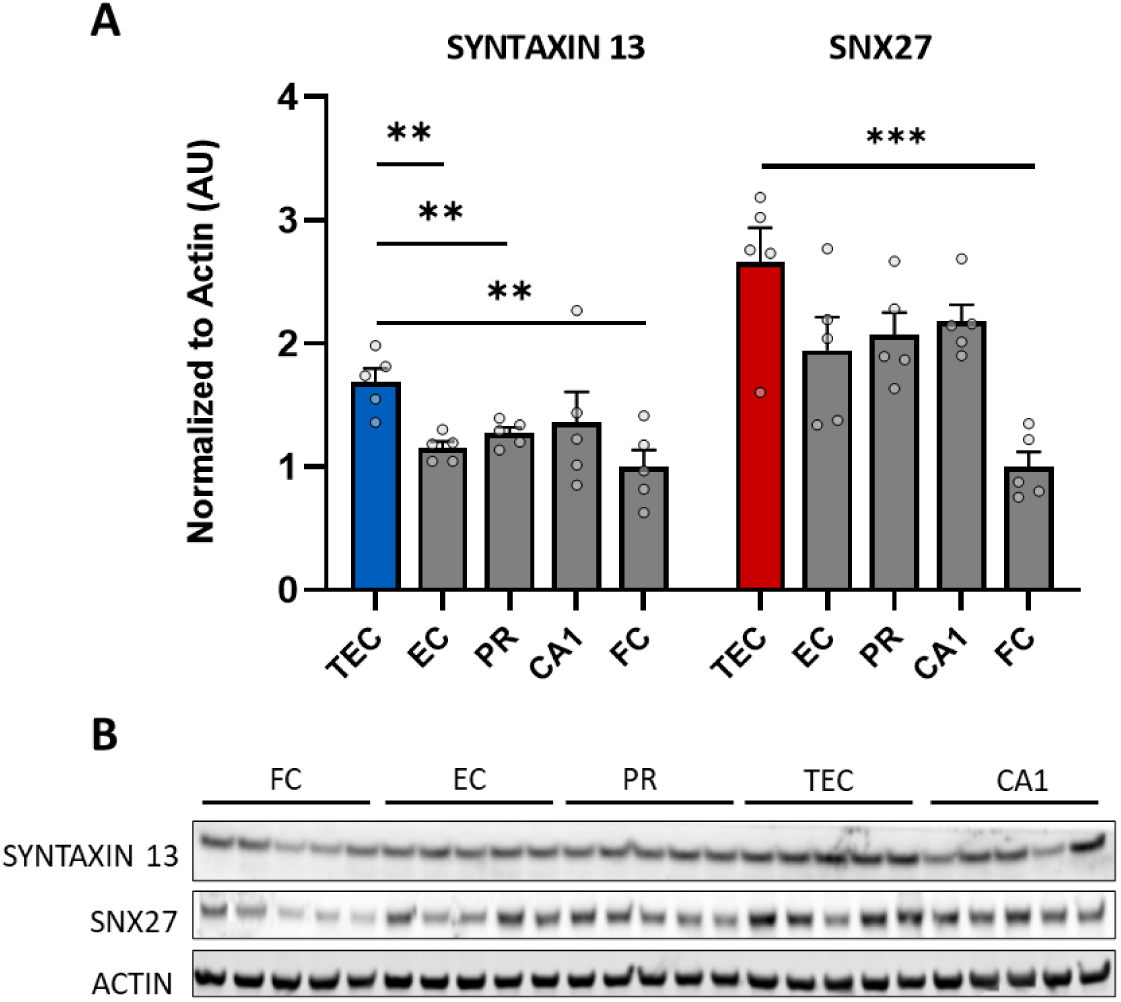
Markers of endosomal recycling in the trans-entorhinal cortex and neighboring brain regions. Compared to neighboring regions, the most reliable differential trans-entorhinal cortex enrichment was found for the recycling-associated SNX27. Syntaxin13 also showed the highest expression in the trans-entorhinal cortex but was only reliably different from the frontal cortex. Data are mean ± SEM and genotypes are compared by two-tailed unpaired Student’s t-test. (n = 5) *\*\*P < 0.01; ***P < 0.001*.

These results offer evidence in support of previous conclusions that, owing to its network properties, TEC neurons differentially engage in endosomal recycling^47^.

## DISCUSSION

By showing that SORL1 deficient mice, modelling causative SORL1 mutations, largely phenocopy the pattern of age-dependent atrophy in AD centered on the TEC, and that atrophy is prevented by VPS26b overexpression and SORL1 restoration, our findings support the hypothesis that disrupting SORL1-retromer endosomal recycling can drive AD’s neurodegenerative process.

The differential vulnerability of TEC neurons to pathway dysfunction was established in a previous study with VPS26b deficient mice^5^, which is explained by a unique network property of the TEC neurons^47^ that function as a cortical hub receiving the highest level of input from across the cortex^46,47^. The prefrontal cortex also has hub-like properties^48^, which might explain why it emerged as the secondary region sensitive to SORL1 deficiency. While network properties can explain the spatial phenocopy to AD, what about the observation that SORL1 deficiency phenocopies the temporal profile of AD, where neurodegeneration is age-dependent and slowly progressive? This question is one of the most important unresolved questions in the field, where even monogenic causal mutations in AD typically manifest in midlife. Our results suggest interlinked mechanisms that can address this temporal question.

First, SORL1 deficiency was found to lead to an age-dependent lowering of SORL1 levels, a process rescued by VPS26b overexpression. In addition, only when reaching a critical threshold is SORL1 deficiency associated with neurodegeneration. This age-dependency agrees with how SORL1 is not only a pathway regulator, but being an endosomal membrane protein, it itself is recycled away from degradation, as shown for example by VPS26b depletion-repletion studies in neuronal culture^5^. Thus, SORL1 deficiency or other pathogenic genetic SORL1 defects that prevent it from interacting with retromer^49,50^ will give rise to a feedback mechanism that can drive down SORL1 with advancing age.

The proteomic analyses in SORL1 deficient mice suggest a second mechanism that can impose a slow and chronically progressive neurodegenerative process. The proteomic profile is primarily characterized, not by an increase in potentially neurotoxic proteins, but rather by the loss of neuroprotective proteins that suppress cell death. Future studies are required to clarify the link between these secondary deficiencies and defective endosomal recycling, but it is plausibly related to the pathway’s function in maintaining the health of lysosomes, autophagosomes, mitochondria, and mitophagy^13–27^. It is because of this function that the pathway has been assigned a “neuroprotective” role^20,51^, and the loss of cell death suppressors we observe here broadens this term to include protection against cell death. Rather than acutely driving widespread and synchronized neuronal death, loss of neuroprotection will render neurons stochastically vulnerable to metabolic stressors, resulting in a slow and asynchronous pattern of neuronal loss. When concatenated with age-dependent worsening of SORL1, these mechanisms can account for why patients with causative loss-of-function SORL1 mutations only clinically manifest in midlife. These mechanisms may also generalize beyond SORL1. For example, recent studies suggest that AD-associated mutations in APP and the PSENs can secondarily lead to retromer deficiency^52,53^, and so the mechanisms might extend to incorporate other monogenic causes of AD.

More importantly, this mechanism might generalize to the common, sporadic, form of AD. Besides the fact that SORL1 is one of the most frequent risk genes found in sporadic AD^54^, it was in sporadic patients that the SORL1-retromer pathway was first implicated in AD by profiling the entorhinal cortex^55^ and a biomarker of pathway dysfunction was found in the cerebrospinal fluid of sporadic patients^56^. By probing the atrophic TEC more specifically, a co-deficiency in SORL1 and VPS26b was observed^5^, and we now find that it is also deficient in NLRX1, identified in the TEC of SORL1 deficient mice.

Besides genetic drivers and co-morbidities like type II diabetes and obesity^57^, recent studies have suggested a positive feedback relationship between pathway dysfunction and the multiple proteinopathies that typifies sporadic AD -- amyloid aggregates, tau aggregates, and TDP-43 aggregates^52,58–60^. When taken together with the fact that mice do not develop these proteinopathies unless engineered to express relevant human proteins, this raises the possibility that disruptions to SORL1-retromer endosomal recycling might represent a final common ‘neurodegeneration pathway’ in AD (Supplementary Figure 3), one that we show here can be potentially targeted therapeutically .

In summary, besides confirming the hypothesis that defects in SORL1-retromer endosomal recycling can drive AD’s neurodegenerative process, the specific details of our findings give rise to testable mechanisms that can explain the age-dependency and slow neurodegenerative time-course of SORL1 defects and potentially in AD more broadly. Additionally, the results add to the convergence of genetics with studies in model systems and patient brains in support of the conclusion that pathway disruptions can be considered causative in AD^12^. More importantly for patients, our results justify the development of drugs that by increasing endosomal SORL1-retromer hold promise in ameliorating or even preventing the disease’s neurodegenerative process, the ultimate cause of its morbidity and mortality.

## METHODS

### In vivo imaging

#### MRI acquisition

High-resolution structural MRI was performed in vivo using a dedicated 9.4 T Bruker BioSpec small-animal scanner equipped with a mouse brain MRI CryoProbe (Bruker BioSpin, Germany). Mice were maintained under isoflurane anesthesia, with body temperature maintained at approximately 37°C and respiratory rate monitored and maintained between 40 and 60 breaths/min throughout imaging. T2-weighted anatomical images were acquired using a rapid acquisition with relaxation enhancement (RARE) sequence with a repetition time (TR) of 2,500 ms, an effective echo time (TE) of 45 ms, an in-plane resolution of 60 × 60 μm², and a slice thickness of 250 μm, yielding a voxel size of 60 × 60 × 250 μm³. The acquisition protocol followed previously established high-field mouse MRI methods^5,39,61^.

#### Image preprocessing and tensor-based morphometry

A study-specific group-wise anatomical template was constructed using Advanced Normalization Tools (ANTs). Brain-extracted T2-weighted images were first linearly aligned and subsequently registered to the template using symmetric normalization (SyN) nonlinear diffeomorphic registration, establishing voxel-wise anatomical correspondence across mice^62,63^. Tensor-based morphometry (TBM) was performed following previously established mouse MRI analysis methods^39^. Voxel-wise Jacobian determinant (JD) maps were calculated from the nonlinear deformation fields to quantify local volumetric changes relative to the study-specific template^61^. Log-transformed Jacobian determinant (log-JD) maps were used for voxel-wise statistical analysis, whereas JD maps were used for regional volume quantification.

#### Voxel-based statistical analysis and cluster definition

Individual log-JD maps were analyzed using a general linear model in Statistical Parametric Mapping (SPM), following the previously established mouse TBM approach^39^. The exploratory analysis compared SORL1^Exon4+/−^ mice with SORL1^Exon4+/+^ controls at 12 months of age. In the principal genotype study, voxel-wise linear regression across SORL1^Exon4+/+^, SORL1^Exon4+/−^, and SORL1^Exon^^4−/−^ mice at 12 months was performed to assess the association between SORL1 gene dose and regional brain volume. To evaluate the potential influence of sex, the voxel-wise regression was repeated with sex included as a covariate. For the intervention study, voxel-wise comparisons were performed between AAV9-empty-vector-treated SORL1^Exon4+/−^ mice and SORL1^Exon4+/+^ controls, between AAV9-VPS26b-treated SORL1^Exon4+/−^ mice and SORL1^Exon4+/+^ controls, and directly between the two treatment groups. A voxel-wise cluster-forming threshold of *P* < 0.005 was applied, and suprathreshold voxels were grouped into spatially contiguous clusters. Cluster-level correction for multiple comparisons was performed using AFNI-AlphaSim Monte Carlo simulation with 5,000 iterations. Clusters meeting the simulation-derived extent criterion corresponding to a cluster-wise corrected *P* < 0.05 were retained^39^. The significant cortical clusters identified in the principal genotype study were anatomically localized to define the trans-entorhinal cortex (TEC) and anterior prefrontal cortex (PFC) regions of interest. TEC localization was cross-referenced to the Allen Mouse Brain Atlas at the border between the lateral entorhinal and perirhinal cortices, consistent with the anatomical definition reported previously^5^.

#### Regional analyses across age and intervention groups

The TEC and PFC regions of interest defined from the corrected 12-month genotype analysis were held fixed for regional analyses of SORL1^Exon4+/+^, SORL1^Exon4+/−^, and SORL1^Exon^^4−/−^ mice at 3, approximately 6, and 12 months of age. Regional volume was quantified for each scan by summing the JD values across all voxels within each ROI, following the previously established approach^39^. These measurements were used to characterize age- and genotype-dependent regional volume changes, with normalized regional volumes presented in Figure 2C. The same TEC region of interest was carried forward for regional evaluation of the 12-month intervention cohort, including AAV9-VPS26b-treated SORL1^Exon4+/−^ mice, AAV9-empty-vector-treated SORL1^Exon4+/−^ mice, and SORL1^Exon4+/+^ controls.

### SORL1^Exon^^4-/-^, SORL1^Exon4+/-^ and SORL1^Exon4+/+^ mice

Mice were generated and described previously^37,38^. Breeding pairs were provided by Dr. Olav Andersen at Aarhus University Denmark.

### AAV9 production

AAV preps were produced at Vector Biolabs.

### Western blots (Immunoblotting)

Proteins from mouse brain were isolated as described previously^32,64^. Lysates from the samples were run on NuPAGE® Bis-Tris 4-12% gels, transferred onto nitrocellulose membranes using iblot and were probed with antibodies. Primary antibodies targeting the following proteins were used for probing; VPS26b (NBP1-92575, Novus, 1:500) and β-actin (ab6276, Abcam), SORL1 (anti-LR11 BD-611861, BD bioscience 1:1k), Synaptophysin (101 002 SYSY 1:2k), Syntaxin 13 (110132 SYSY 1:1k), SNX27 (NBP1-45283 Novus Bio 1:1k), NLRX1 (Cat #:17215-1-AP Protein Tech 1:1k). The primary antibodies were selected based on extensive citations by previous studies; minimal non-specific signal and validation by KO testing. IRDye® 800 or 680 bound antibodies (LI-COR), or HRP bound antibodies against the host were used as secondary with dilutions of 1:10k for 800CW, 1:15k for 680RD, 1:25k for 680LT antibodies, and 1:2k for HRP antibodies. Western blots were scanned using the Odyssey imaging system as described previously^65^.

### Stereotactic injection of AAV9 in mouse

All animal procedures were approved by Columbia University IACUC. AAV9 viral particles were injected into the Inferior Horn of Lateral Ventricle (IHLV) region of the mouse brain. IHLV region was chosen because of association of this region with TEC, where the atrophy was detected using MRI. Briefly, the surgical procedure was carried out in an aseptic environment. Animals were anesthetized using ketamine / xylazine. Fur was clipped and the site cleaned with betadine scrub followed by 70% ethanol. Eyes were lubricated with an application of sterile, bland ophthalmic ointment. Preemptive analgesia was induced by carprofen and marcaine injections. Mouse was placed on infra-red warming pad in the stereotaxic frame connected to an automatic injection pump. Surgeon wore proper protective equipment and sterile gloves. Using sterile scissors, skin was cut, and bone exposed. A burr hole was placed in the skull at the experimental coordinates calculated using landmarks (bregma and lambda). The dura mater was pierced using a sterile needle and the syringe needle lowered to the desired z coordinate.

AAV9 virus was injected through a 33g needle attached to a Hamilton neuros syringe at a rate of 1 μL/min over a period of 10 min (10 μL volume). Optimal dose was identified using a dose response ranging from 4E+9 to 1E+11 vector genomes. The needle was retracted 5 min after the end of injection. Surgical wound was closed/sutured by a combination of skin glue and sterile stitches. Post-surgery, 1 mL of normal saline was injected via intraperitoneal route, to minimize dehydration. Infra-red warming pad was used during post-op recovery. Carprofen injection was given once daily for 3 days post-op. Animals were monitored continuously until ambulating and once daily for the first 3 days and then twice a week until day 14 post-op. Sutures were removed on day 10-14 post-op. Animals were injected at 3 to 3.5 months of age, and the brains were harvested 12-15 months post-surgery, for analysis.

### Perfusion and tissue processing

After the survival period, the animals were perfused with saline, the brains extracted, and the site of injection was microdissected out of the brain for biochemical and proteomics analysis.

### Immunohistochemistry

For immunohistochemistry, mice were transcardially perfused with 4% paraformaldehyde. Brains were removed and post-fixed with 4% paraformaldehyde for 24 hours at 4°C, followed by cryoprotection with 30% sucrose for 72 hours at 4°C. Brains were embedded in Optimal Cutting Temperature embedding medium (Tissue-Tek) and stored at -80°C. Frozen brains were sectioned on a cryostat into 20µm thick sections, and mounted. Coronal sections were chosen to include the TEC.

Frozen mouse brain sections were washed in 1X phosphate buffered saline (PBS) 3 times for 5 minutes, permeabilized with 0.1% Triton X-100 for 20 minutes, and blocked overnight in blocking buffer (1% bovine serum albumin (BSA), 10% Normal Goat Serum (NGS)) at 4°C. Neurons were detected using an antibody for NeuN (Cell Signaling Technology #D4G4O, 1:250)) prepared in blocking buffer, incubated overnight at 4°C. Afterwards, sections were incubated overnight in a secondary antibody (Invitrogen #A-11012, 1:2000), and stained with the nuclear stain DAPI (concentration?). To reduce autofluorescence, sections were incubated with 0.1% Sudan Black B in 70% ethanol. Sections were then washed with 1X PBS and mounted with Fluoromount-G.

Sections were imaged on an Andor BC-43 confocal microscope, using the rhinal fissure and external capsule as anatomical landmarks to identify the TEC. Images were acquired using a 60x objective and stitched, covering a region of approximately 3mm^2^. Images were segmented using Ilastik^66^, and neurons were counted using Fiji. NeuN and DAPI-positive overlap in approximately 1mm length of cortex, including all cortical layers were counted. All analyses were conducted with the investigator blinded to the treatment group.

### Proteomics data acquisition

Mouse brain tissues were resuspended in 200 µL of solubilization buffer (1% SDC, 2% SDS, 100 mM Tris pH 8.0, and protease inhibitors) and homogenized in Preomics’ BeatBox at hard settings for 10 minutes. The protein lysates were centrifuged at 14,000 rpm for 5 minutes, and the supernatants were transferred to new 1.5 mL Eppendorf tubes. Protein concentrations were determined using the Pierce BCA Protein Assay Kit, and 10 µg of protein from each lysate was used for further processing. Samples were boiled for 20 minutes at 60°C with agitation (1000 rpm). Protein reduction and alkylation of cysteine residues were performed using 10 mM tris(2-carboxyethyl) phosphine (TCEP) and 40 mM chloroacetamide (CAA) at 45°C for 15 minutes. Following sonication in a water bath and cooling to room temperature, proteins were digested overnight with a LysC/trypsin mix at a 1:50 enzyme-to-protein ratio at 37°C and 1400 rpm. The resulting peptides were acidified to 1% trifluoroacetic acid (TFA) and desalted using Empore SDB-RPS StageTips. Eluted peptides were dried using a SpeedVac concentrator and resuspended in 3% acetonitrile containing 0.1% formic acid. Peptide concentrations were measured using a NanoDrop spectrophotometer. For LC-MS/MS analysis, 130 ng of peptides per sample were analyzed, and 200 ng of each sample were used for diaPASEF acquisition on a timsTOF Pro 2.

Liquid Chromatography–Tandem Mass Spectrometry (LC–MS/MS): Peptides were separated over a 60 min gradient at a flow rate of 400 nl/min on a reverse-phase C18 column with an integrated CaptiveSpray emitter (25 cm × 75 µm, 1.6 µm; IonOpticks). Mobile phase A consisted of 0.1% formic acid in water, and mobile phase B consisted of 0.1% formic acid in acetonitrile. The gradient was programmed as follows: 2% to 25% B over 45 min, 25% to 40% B over 5 min, then increased to 95% B before column re-equilibration. The timsTOF Pro2 was operated in diaPASEF mode^67^. Data were acquired using 60 variable isolation windows covering m/z 350–1,250. Collision energy was linearly ramped with ion mobility, ranging from 59 eV at 1/K₀ = 1.45 Vs cm⁻² to 20 eV at 1/K₀ = 0.6 Vs cm⁻².

### Proteomic Data Processing and Statistical Analysis

Quantitative mass spectrometry was analyzed using the DIA-Analyst web application. Intensity values were *log*_2_-transformed and normalized. Differential protein expression between SORL1 deficient mice and wildtype littermates was evaluated using moderation via *t*-statistics. Multiple testing correction was performed using t statistic-based False Discovery Rate (FDR) correction implemented in DIA-Analyst. Proteins were considered significantly differentially expressed if they met an adjusted *p*-value threshold of ≤ 0.05 combined with an absolute fold change cutoff of 0.25

## Acknowledgment

This work was partly funded by Retromer Therapeutics and grants from the Alzheimer’s Association, the Taub Institute and the Ludwig Foundation and by the NIH: P30 AG066462 (Alzheimer’s Disease Research Center) and P01AG09268 (SORL1 in Alzheimer’s disease pathogenesis and pathophysiology). We thank Dr. Olav Andersen, Aarhus University Denmark, for providing the mice.

## Disclosures

SAS is on the Scientific Advisory Board of Graviton Biosciences and Leal therapeutics

---

**Supplementary Figure 1.**
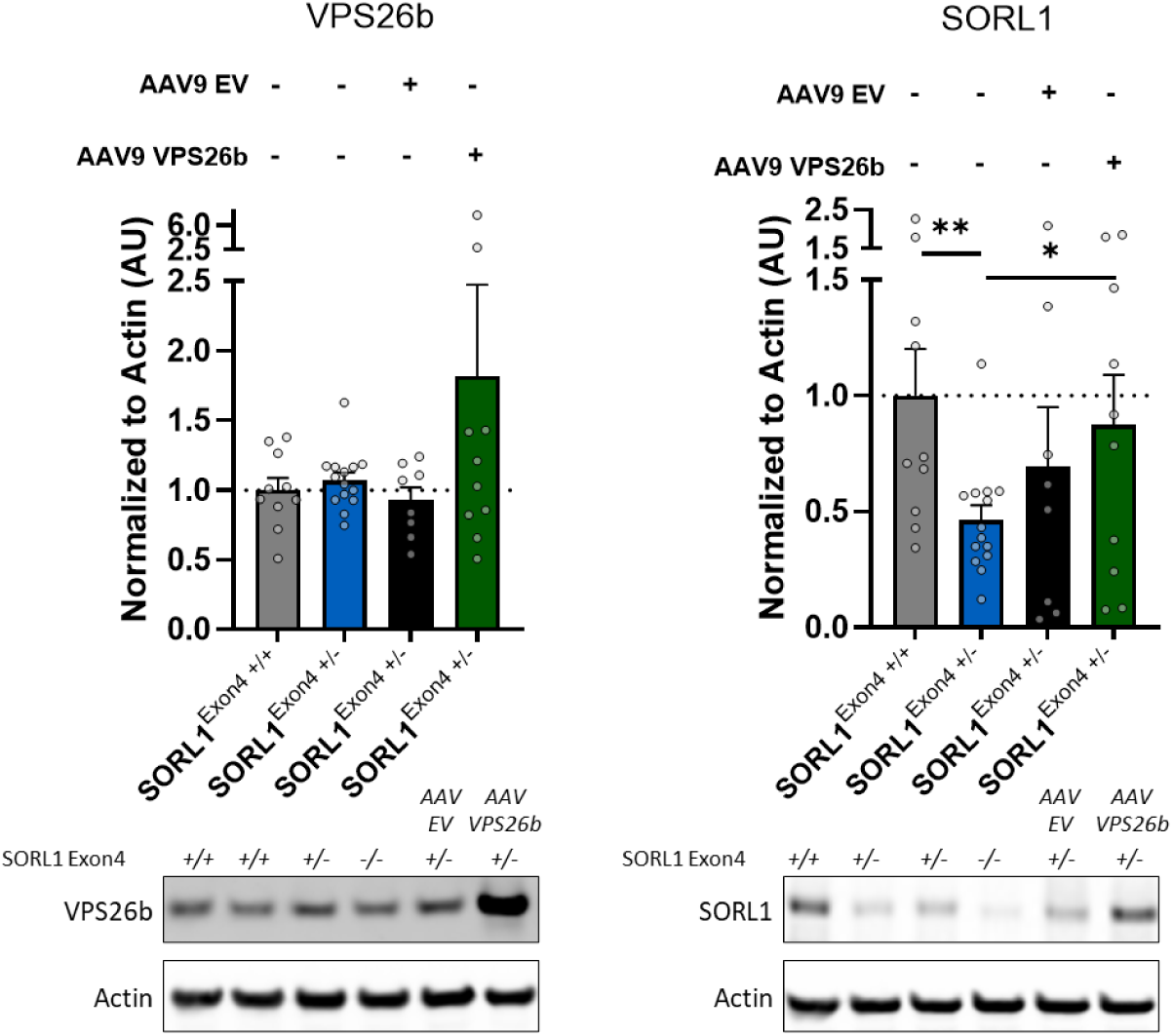
VPS26b and SORL1 protein levels after AAV9 treatment. 12-month-old trans-entorhinal cortex tissue from the treated mice confirmed that the only treatment with AAV9 VPS26b results in increased VPS26b expression. This VPS26b overexpression was associated with increased SORL1 levels, effectively normalizing SORL1 deficiency. AAV9 Empty Vector (EV) also resulted in a non-significant increase in SORL1 levels, however there was no increase in VPS26b levels. Data are mean ± SEM and genotypes are compared by two-tailed unpaired Student’s t-test. (n = 10, 14, 8, 10)*\*P < 0.05, **P < 0.01*.

**Supplementary Figure 2.**
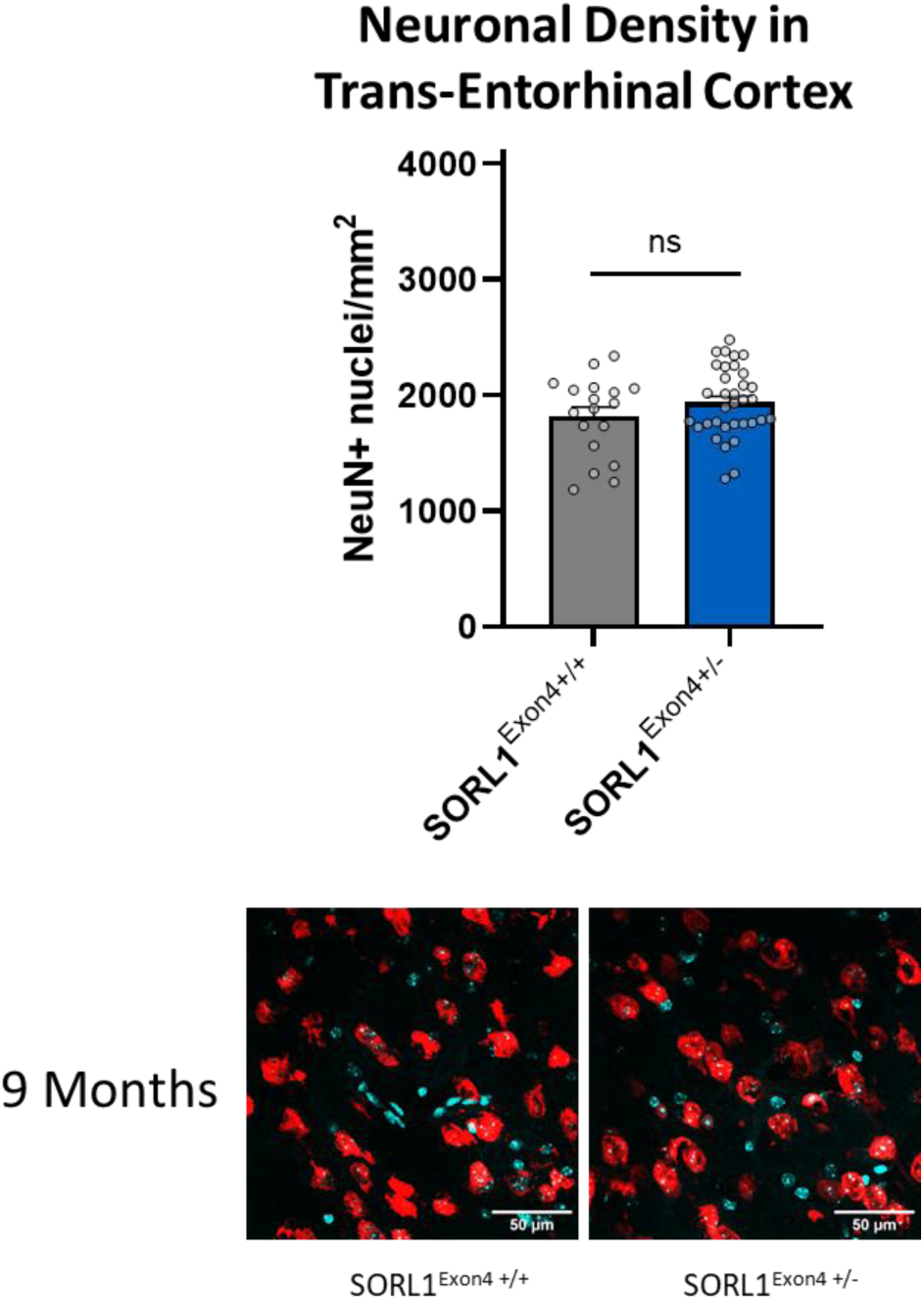
Neuronal Density in Trans-Entorhinal Cortex at 9 Months. Confocal microscopy, in trans-entorhinal cortex of 9 month old mice. The bar graph shows the quantification of NeuN+ nuclei per mm2 of cortex. Data are mean ± SEM and genotypes are compared by Kruskal-Wallis test. *\*P < 0.05, **P < 0.01; ***P < 0.001, ns = not significant.* Representative immunofluorescent images of NeuN (red) and DAPI (blue) stained in the trans-entorhinal cortex of aged SORL1-deficient mice (N=5-8 mice per genotype). Scale bar, 50µm.

**Supplementary Figure 3.**
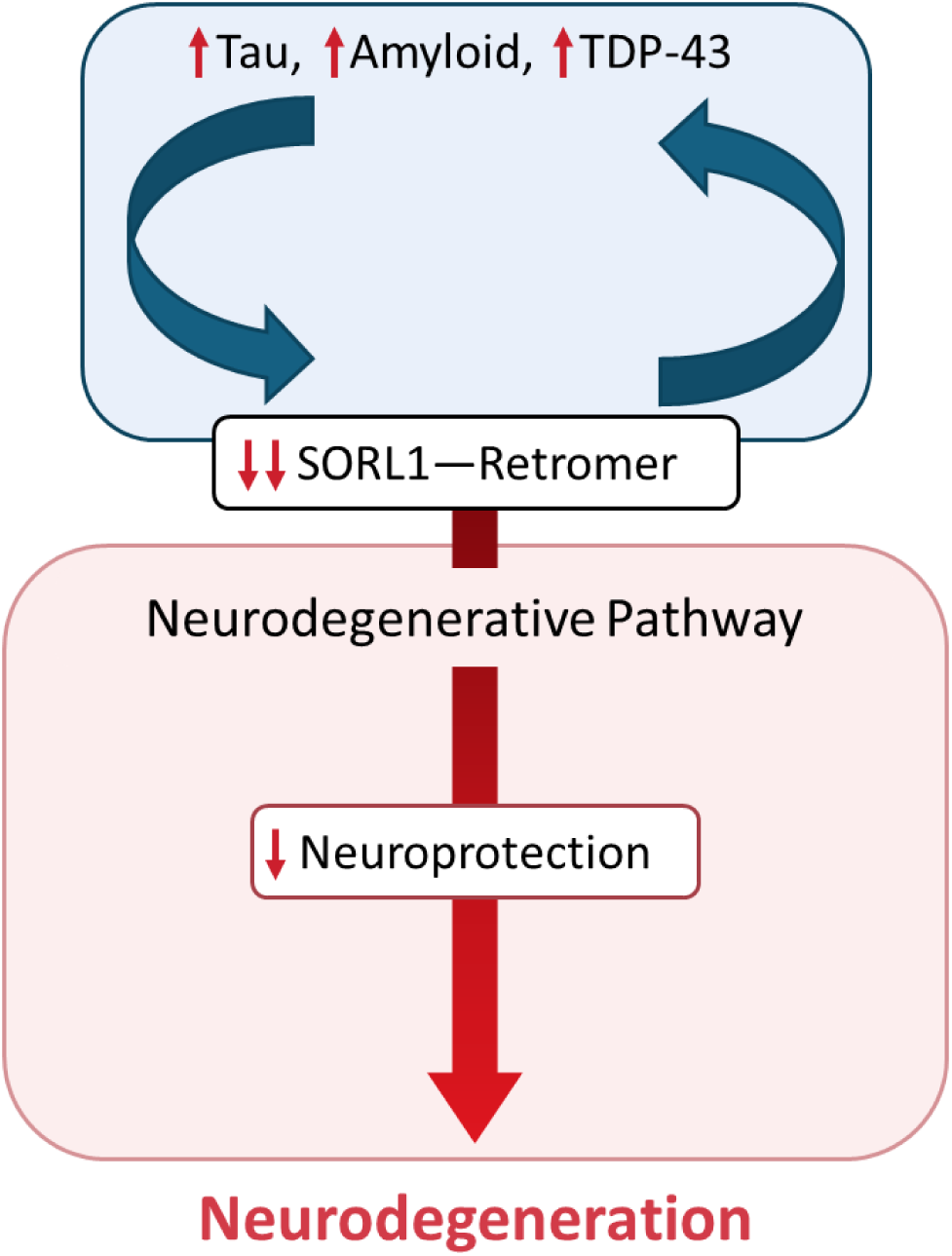
A proposed common neurodegenerative pathway in Alzheimer’s disease. A positive feedback relationship exists between SORL1-retromer dysfunction and AD-associated proteinopathies. When SORL1-retromer dysfunction achieves a critical threshold, it induces a loss in neuroprotection, which can trigger the slow neurodegenerative process that typifies the disease.

